# Metacaspases mediate thermotolerance of diatoms following marine heat waves

**DOI:** 10.1101/2025.06.09.658587

**Authors:** Mai Sadeh, Daniella Schatz, Shifra Ben-Dror, Avia Mizrachi, Shiri Graff van Creveld, Amichai Zafrin, Assaf Vardi

## Abstract

- Marine heat waves (MHW) are extreme climate events characterized by elevated ocean temperatures lasting from days to months and spanning thousands of kilometers. Recent predictions show an increment in MHW frequency, duration and intensity worldwide along with climate change. Nevertheless, little is known about MHW impacts on marine microbial life and, specifically, on algal blooms. Recent studies in yeast and green algae suggested that thermotolerance is mediated by metacaspases (MC), cysteine proteases, structurally similar to caspases known to mediate programmed cell death.
- Initially we created a heatwave model of 72-hours that exposes the model diatom *Phaeodactylum tricornutum* to elevated temperature with a recovery phase that enables to capture the mechanisms underpinning such acclimation. We generated triple knock-out mutants of the PtMCA genes and identified a vital role for MC in heat-stress acclimation.
- PtMCA triple mutants exhibited increased sensitivity to the heatwave treatment, which induced cell death that peaked days after returning to initial temperatures. We further revealed that heatwave treatment induced accumulation of reactive oxygen species and the PtMCA mutants were hypersensitive to oxidative stress as compared to WT cells.
- We propose that metacaspases have a pivotal role in diatom’s acclimation to elevated temperatures a trait vital for algal survival considering climate change.

## Introduction

Ocean temperatures are rising due to global climate change, and this warming is often accompanied by extreme fluctuations known as marine heatwaves (MHWs) (Oliver *et al*., 2018; Intergovernmental Panel On Climate Change (Ipcc), 2023). There are several ways to define MHWs, but most definitions describe MHWs as extreme rises in ocean temperature above multi-annual averages for an extended period ranging between days and months. The severity of the MHW depends both on the temperature and the duration, both of which are increasing in recent years, and are predicted to further increase in the future (Hobday *et al*., 2016). Some MHW s can expand over thousands of kilometers. MHWs can be region-specific, though extreme MHWs are usually linked to large-scale climate changes (Holbrook *et al*., 2019). The effect of MHWs on marine biology has been studied mainly in the context of coral reef bleaching and mass mortality of benthic communities (Garrabou *et al*., 2009; Genin *et al*., 2020; Kochman-Gino & Fine, 2023), yet little is known about their effect on marine alga. Heat stress anomalies were shown to cause various effects during phytoplankton blooms (Kim *et al*., 2024) and were shown to inhibit the annual phytoplankton bloom in the north-west Mediterranean sea (Rosalia Santoleri, 2024).

Diatoms are among the most diverse and geographically distributed phytoplankton groups. They have a major contribution to global primary production that can reach up to 20% (Armbrust, 2009; Pierella Karlusich *et al*., 2024). The response of diatoms to heat stress was studied by different approaches and experimental setups, demonstrating that diatoms can adapt to diverse temperature niches, yet their cellular response to heatwaves is underexplored. In the centric diatom *Skeletonema marinoi,* comparison of resurrected strains from the past 60 years demonstrated that modern strains shifted their optimal growth temperature by 1°C compared to older strains (Hattich *et al*., 2024). The Southern Ocean diatom *Actinocyclus actinochilus*, had mixed effects after exposure to heatwaves (HW); longer HW treatments resulted in higher cell death, though higher temperatures during the HW resulted in a higher optimal growth temperature (Samuels *et al*., 2021). Adaptive laboratory evolution approaches were also used to expand the understanding of HW effects. One outcome of severe warming treatment of 28°C in *Skeletonema dohrnii* was the significant reduction in the average number of nucleotide differences per site in the genome, also known as nucleotide diversity. Warming treatment resulted in reduction of 10% in nucleotide diversity (compared to populations in ambient temperature) that may harm potential of individual cells and populations to adapt to rising temperatures (Cheng *et al*., 2024).

Transcriptomics approaches revealed differentially expressed genes and metabolic pathways associated with growth under optimal and sub optimal temperatures in *Chaetoceros* strains adapted to different temperatures (Liang *et al*., 2019). Cold-adapted strains invested more in RNA transporters, translation factors and tRNA synthesis while warm adapted strains divested of ribosomes to reduce unfolded proteins and ER stress. It was suggested that diatoms developed temperature niches through changes in baseline expression in key pathways including lipid metabolism, protein folding and degradation, and cell death (Liang *et al*., 2019). In the model diatom *Phaeodactylum tricornutum,* transcriptome analysis of cells exposed to elevated temperatures showed shutdown of pathways related to photosynthesis. In contrast nitrogen metabolism was activated to accumulate internal storages and was also suggested to be correlated with cell size increment that reduces susceptibility to heat stress (Hong *et al*., 2023, 2025).

Mechanisms to cope with heat stress were shown in depth in other model systems such as yeast (Verghese *et al*., 2012) and *Chlamydomonas* (Zhang *et al*., 2022). In both systems recent findings highlighted that metacaspases (MC) may serve as key proteins involved in acclimation to heat-stress. This is in contrast to the classic view of the role of MCs in programmed cell death (PCD) (Lee *et al*., 2010; Zou et al., 2023).

Traditionally the role of MCs in plants and unicellular organisms was thought to be linked to cell death due to their structure similarity to caspases, cysteine proteases that are canonical cell death executors in mammalian systems (Kaczanowski *et al*., 2011). Since unicellular organisms and plants lack caspases, it was suggested that metacaspases are involved in PCD. However, in various organisms such as yeast (Lee *et al*., 2010), green algae (Zou et al., 2023) and higher plants (Ruiz-Solaní *et al*., 2023), MCs were shown to have different roles in the cell, such as maintaining cellular homeostasis and participating in cell growth (Tsiatsiani *et al*., 2011; Ruiz-Solaní *et al*., 2023). In organisms with several MCs, such as in the model plant *A. thaliana*, which has 9 MC genes, each MC has a different role. Some were demonstrated to have cell death roles while others were crucial for defense against pathogens (Coll *et al*., 2010; Watanabe & Lam, 2011; Salguero-Linares *et al*., 2025). Furthermore, often the same MC could have different functions that depend on the cellular context. For example, the yeast Mca1 switches from protease function that promotes cell death to co-chaperone activity that delays aging, through the binding of calmodulin to its pro-domain (Eisele-Bürger *et al*., 2023). Previous work shed light on the wide abundance of MCs in diatoms and revealed that *P. tricornutum* has 5 MCs, two of which (PtMCA-a and PtMCA-b) lack the p10 domain and are considered pseudogenes. The remaining three MCs are type III (MCAIII). Of these, PtMCA-IIIc had the highest expression levels, and we previously showed its role in activation of cell death in response to oxidative stress (Graff Van Creveld *et al*., 2021). The abundance of the MCAIII among diatoms was shown and additional biochemical characterization of PtMCA-IIIc revealed the calcium dependency of the MC proteolytic function (Graff Van Creveld *et al*., 2021). The functional role of PtMC-IIIc and the two other expressing MCs (PtMC-IIIa and PtMC-IIIb) in *P. tricornutum* still remains unknown.

In the current study, we aim to investigate the role of metacaspases in driving thermotolerance in marine diatoms. We focus on the stress response and acclimation of *P. tricornutum* to short term thermal anomalies. We developed a lab MHW setup using the diatom model *P. tricornutum* and study physiological dynamics throughout the heatwave. We revealed a novel role for PtMCAs and demonstrate their involvement in thermotolerance using genetic, physiological and biochemical approaches.

## Results

In order to study cellular mechanisms in response to heat stress, we grew *P. tricornutum* cells in a range of temperatures. Typically, *P. tricornutum* is maintained in our lab at 18°C, and has a known growth temperature range between 18 and 22°C. We first tested *P. tricornutum* response to shifts in temperature, measuring growth rates during transition from 18°C to higher temperatures (Fig. 1A). Growth rate decrease from 0.75±0.3 to about 0.55±0.5 µ day^-1^ when *P. tricornutum* was transferred from 18°C to 23°C and 28°C, (Fig. 1A). A negative growth rate, suggestive of cell death was measured following the transition from 18°C to 33 and 35°C. To complement these results, we measured the percentage of dead cells, before (day 0), and 2 days after the temperature shift, finding increase in the percentage of dead following elevating the temperature to 33°C (18% ±2) and 35°C (98% ±0.5) (Fig. 1B). While there is no observed cell death after two days at 30°C or below, most of the population dies after two days at 35°C. At 33°C, there was a partial death of the population that indicates severe stress, though this treatment was not completely lethal to the entire population as in 35°C. By quantifying cell death after two days, we can assess which temperature causes mild stress and which severe stress after a short period of time (Fig. 1B). We identified the temperature of 33°C as the tipping point, above which most of the population dies from heat stress, therefore we chose to continue with transition from 18°C to 33°C as a heat-stress temperature.

**Figure 1.**
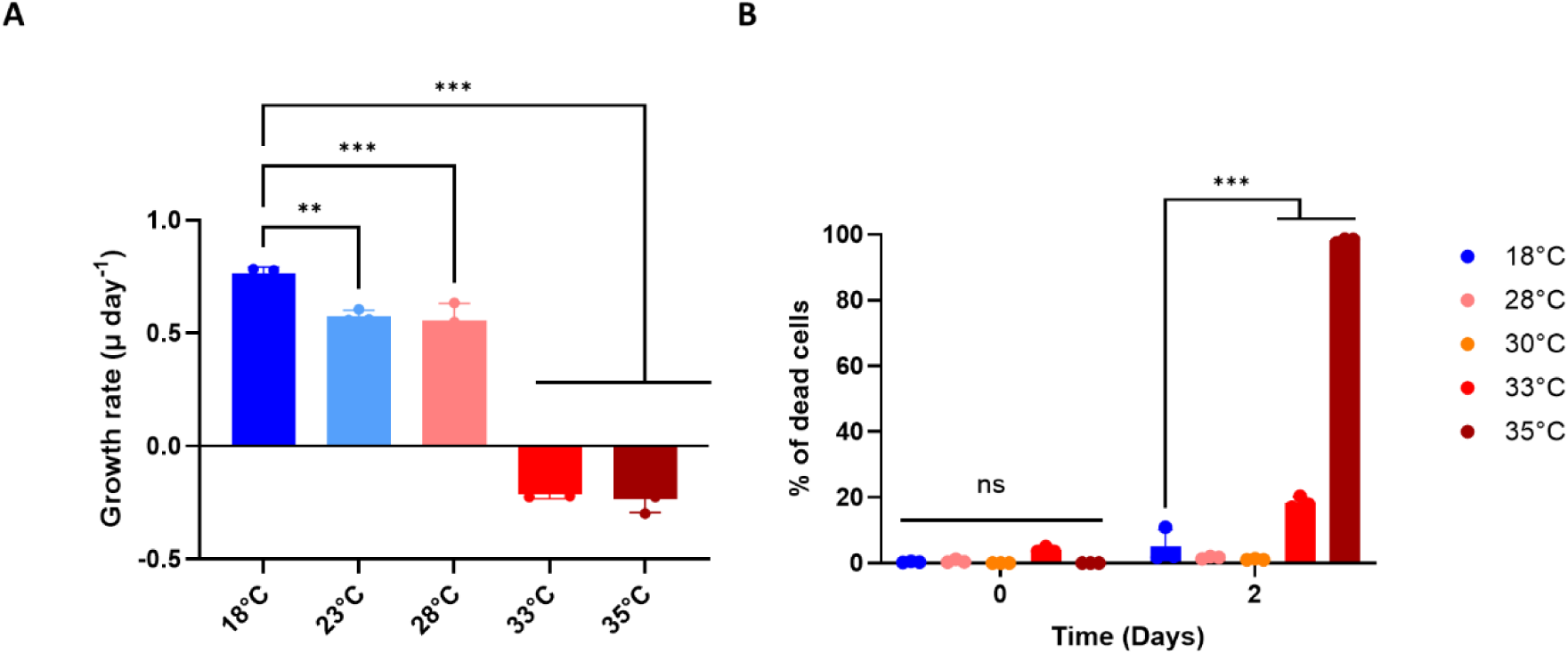
*P. tricornutum* growth rate and cell death in response to elevated temperatures. **(A)** *P. tricornutum* was transferred from its growth temperature of 18°C to 23°C, 28°C, 33°C and 35℃, and cell abundance was measured by flow cytometry. Growth rate was calculated using equation A (see methods). **(B)** Cell death measurements of *P. tricornutum* following transition from 18°C to 28℃,30℃, 33℃ and 35℃ at time zero (before transferring) and after two days using Sytox green staining and flow cytometry. (**A-B**) Values represent the mean ± standard deviation (sd), n=3. *p* values were calculated using 2-way ANOVA compared to 18℃.P values: **<0.02 ***<0.0001. ns= not significant.

Next, we sought to establish a simple lab setup that would not only mimic transient exposure to elevated temperature but will also capture the recovery phase and the mechanisms underpinning such acclimation. To study the response of *P. tricornutum* to MHWs, we designed a HW experimental setup that induces stress and partial mortality by transition from 18℃ to high temperature of 33°C for a defined duration, followed by recovery at the return to basal temperature of 18℃. First, to investigate how different exposure time at elevated temperatures affect *P. tricornutum*, cultures grown at 18°C were transferred to 33°C for 1-3 days then transferred back to 18°C and were monitored for their growth and mortality using flow cytometry (Fig. S1). Cells subjected to 33 °C for 1 day exhibited recovery, manifested by an increase in cell abundance after returning to 18°C, together with decrease in the percent of dead cells at the same time (Fig S1 B-C). Longer exposure to heat stress led to increased mortality and prolonged growth delay (Fig. S1 B-C). Treatments of 24 and 48 hours in 33℃ resulted in a peak of maximum cell death that decreased when cells were transferred back to 18℃ as opposed to 72-hour treatment that resulted in a maximum cell death peak 2 days after transferring the culture back to 18℃. For subsequent experiments, we selected the HW experimental setup of 72 hours at 33°C followed by recovery at 18°C. This setup induced a short-term stress accompanied by gradual induction of cell death that reaches up to ∼55% of the population but also enables a recovery phase, where the percent of dead cells reduced back to the initial values (Fig. 2 and Fig. S1). Intriguingly, cell death peaked 2 days after the return to 18°C, suggesting that a regulated cascade of programmed cell death is activated during the exposure to heat stress and continues after the stress had ended (Fig. 2 A-C and Fig. S1). Growth of the populations arrested after the onset of the heat stress and declined until day 6 (Fig 2A) after which the population started to recover (Fig 2A), 3 days after the HW has ended, when cell death levels started to decrease as well (Fig 2B). Accordingly, photosynthetic efficiency (Fv/Fm) also decreases dramatically from the first day, reaching below 0.1 on day 3, exhibiting recovery from day 6 (Fig 2C). Therefore, we divided the cellular response during the HW to three distinct phases: 1) the heat stress incubation phase (HW), days 1-3, where growth is arrested, and cell death starts to rise gradually; 2) the latent phase (LP), days 3-6, when growth is still arrested and cell death peaks to its maximum; 3) the recovery phase (RP), days 6-9, when growth and photosynthetic efficiency are restored, and cell death in the population declines. Overall, a 3-day heat stress had severe effects on *P. tricornutum* for 3 days after returning to ambient temperature, after which full recovery was observed.

**Figure 2.**
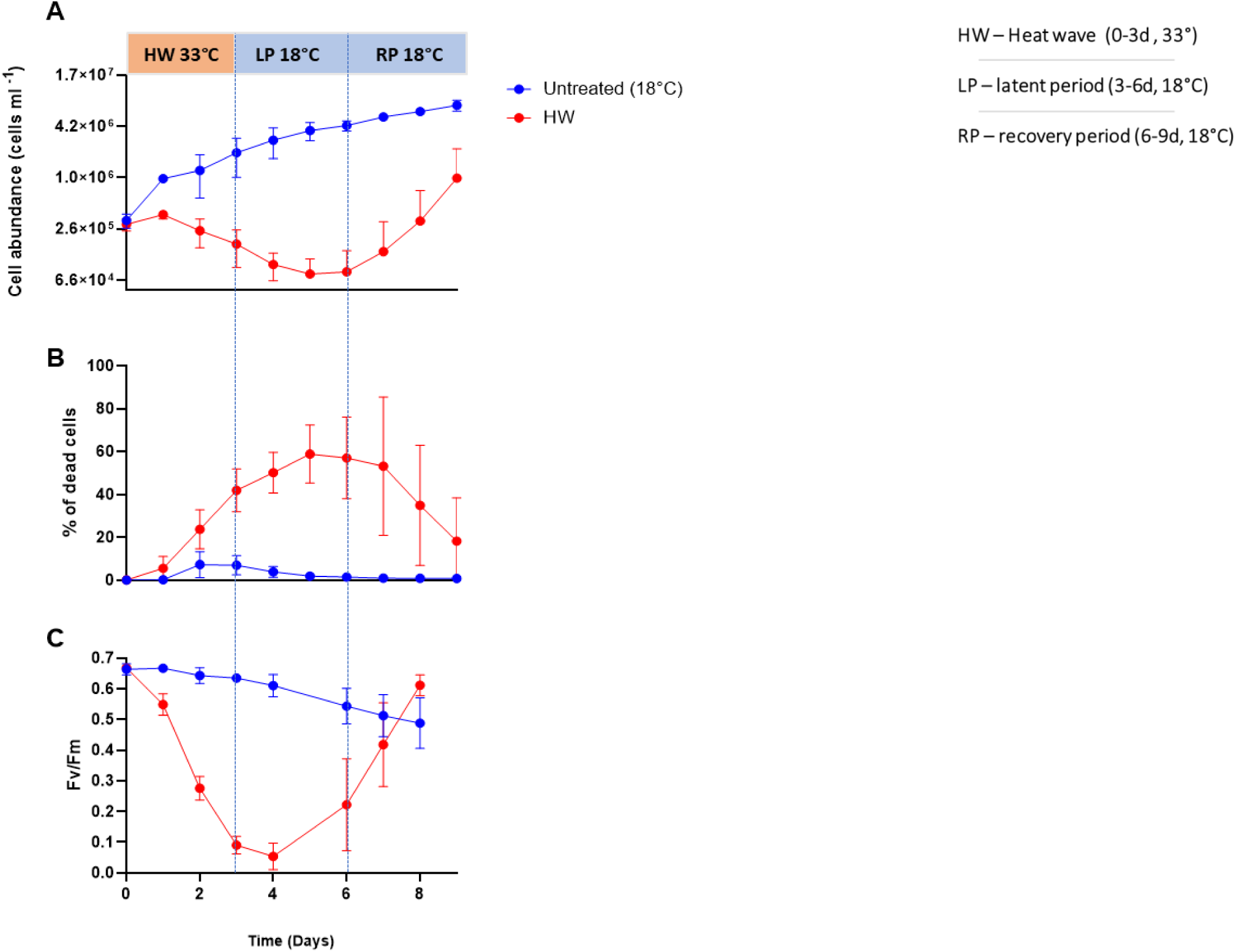
Three phases of physiological responses to HW treatment. *P. tricornutum* cultures were monitored during and after exposure to HW treatment of 72h at 33°C followed by a shift back to 18°C. (**A)** Culture growth, cell counts quantified by flow cytometry. (**B)** Percent of dead cells quantified by flow cytometry using the fluorescent stain Sytox Green. (**C)** Maximal photosynthetic efficiency (Fv/Fm) as measured by Water-PAM. (**A-C**) Blue vertical dashed line represents the separation between the three phases - HW, latent phase (LP) and recovery period (RP). Values represent the mean ± standard deviation (sd), n=3.

Using this experimental setup, we chose to focus on the role of MC in thermotolerance. *P. tricornutum* has 5 MC genes (Fig 3A) (Graff Van Creveld *et al*., 2021). Two of them (PtMCA-a and PtMCA-b) lack the p10 domain and are suspected to be pseudogenes since they are not expressed (Smith *et al*., 2016; Matthijs *et al*., 2017; Graff Van Creveld *et al*., 2021). We previously characterized the proteolytic activity of PtMCA-IIIc and generated single KO lines of PtMCA-IIIc)*ΔMCA-IIIc*) (Graff Van Creveld *et al*., 2021). Since a single deletion of PtMCA-IIIc still resulted in proteolytic activity (Graff Van Creveld *et al*., 2021), we generated double and triple KO lines, thus eliminating all PtMCA-III genes to reveal their full role without potential compensation by other MCs. Using CRISPR-Cas9, we introduced two new mutations to the background strain *ΔMCA-IIIc_1* and disrupted PtMCA-IIIb and PtMCA-IIIa. Some colonies had double KO, creating two lines with double MC KO mutants *ΔMCA-IIIbc_1* and *ΔMCA-IIIbc_2*. Mutants *ΔMCA-IIIbc_1* and *ΔMCA-IIIbc_2* had a frame shift causing a replacement of 155 and 77 amino acids, respectively, in PtMCA-IIIb (Fig S3C). From these colonies two more colonies were isolated that were validated as triple MC KO, *ΔMCA-IIIabc_1* and *ΔMCA-IIabc_2* (see methods). The triple KO strains *ΔMCA-IIIabc_1* has a have deletions of 65 and 3 amino acid deletion in each allele and *ΔMCA-IIIabc_2* has a 68 amino acid deletion in both alleles (Fig S4C).

**Figure 3.**
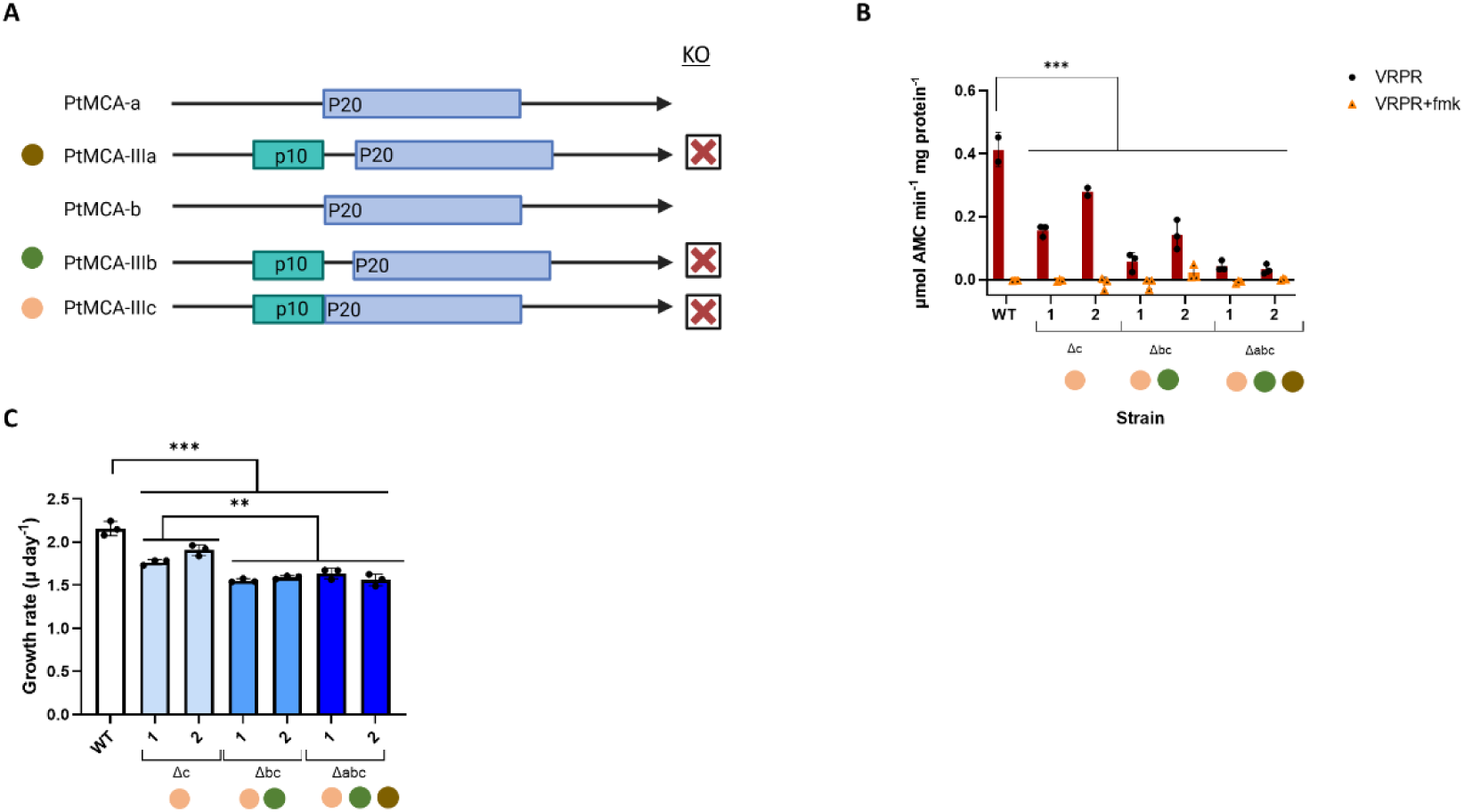
PtMCA-III mutants have impaired MC activity and cell growth. **(A)** Schematic representation of the five PtMC genes in *P. tricornutum*. MC numbers, types, p10 and p20 domain architecture are shown. Red X represents the MC genes that were disrupted in the triple KO mutants. **(B)** Protease activity was measured *in vitro* in extracts from exponentially growing *P. tricornutum* cells. Activity was measured by the release of AMC from peptidyl substrate VRPR-AMC, with and without the inhibitor VRPR-fmk. **(C)** Growth rate of WT versus single, double and triple PtMCA KO mutants. Cells were counted using flow cytometry, and the specific growth rate was calculated as described in the method section**. (B-C)** Values represent the mean ± standard deviation, n=3. Statistical significance was calculated by one-way ANOVA compared to WT. Asterisks represent P-values: **<0.002, *** ≤ 0.0001.

We assessed MC activity using Ac-VRPR-AMC, a fluorogenic metacaspase tetrapeptide substrate that was shown as an optimal substrate for MCs in plants and phytoplankton (Vercammen *et al*., 2006; Spungin & Berman-Frank, 2019). A single KO mutant of only PtMCA-IIIc indeed decreased MC proteolytic activity *in vitro* from a mean of 0.4 in the WT to 0.2±0.5 µmol AMC min^-1^ mg protein^-1^ (Fig 3B), and double mutants *ΔMCA-IIIbc_1* and *ΔMCA-IIIbc_2* further decreased MC proteolytic activity to 0.1±0.5 µmol AMC min^-1^ mg protein^-1^ (Fig 3B). However, in the triple KO mutants MC activity was nearly abolished to 0.03 µmol AMC min^-1^ mg protein^-1^ (Fig 3B). This may imply redundancy or compensation mechanisms when only one or two MCs are non-functional (Fig 3B). In addition, initial characterization of the growth rates of the single, double and triple ptMCs mutant demonstrated their slower growth compared to WT cultures, hinting to their vital role (Fig 3C). Similar to the proteolytic activity, growth rate is also decreased when more MC are KO. Single KO lines also have a significantly faster growth rate compared to the double and triple KO lines, implying that when one MC gene is knocked out there is compensation by the two others (Fig. 3C). Due to their significantly low activity in the triple KO, these lines were selected as a reliable model for studying the role of ptMCA-IIIs in the response to heat stress in *P. tricornutum*.

We characterized the thermal response curve of the triple MC KO and compared it to WT. When grown in elevated temperatures MC triple KO lines had lower growth rates. At high temperatures such as 28°C, growth was almost completely inhibited in the triple KO mutants while the WT is still able to grow with only minimal growth inhibition (Fig 4A). In addition, the critical temperature above which cell death was induced in the mutant lines was lower than that in the WT. WT cells initiate cell death in 33℃, while in the triple KO exposed to 28°C, the percent of cell death in the population reached 40%, suggesting that the tipping point is 5℃ below that for the WT (Fig 4A-B). While a slight and significant difference in growth rate was detected under 18℃ (Fig. 3C, 4C), the HW treatment exacerbated the differences between the WT and the triple KO lines. Throughout the HW exposure the triple mutants had higher fraction of cell death and slower growth rates (Fig 4C-D). Following the HW treatment, the triple mutants also reached the recovery phase in later timepoints. The highest mean percent of dead cells in WT was on day 6 (55%) and for the MC mutants the highest mean was on day 7 (88%) (Fig. 4C-D). Thermal sensitivity is significant only when all three PtMCA-III genes were knocked-out, as single and double mutant strains do not have higher death rates following exposure to HW (Fig S6). This may be explained by the redundant activity in single and double PtMC KO lines (Fig 3B), or by the importance of PtMCA-IIIa specifically in heat-stress response. Based on the hyper-sensitivity of the triple PtMCA mutants to HW, we propose that MCs have a vital role associated with acclimation to thermal stress. To examine the specificity of PtMCAs to elevated temperature, we exposed the cells to two different infochemicals produced by diatoms, that are known to induce PCD, cyanogen bromide (BrCN) (Vanelslander *et al*., 2012; Graff Van Creveld *et al*., 2021) and the oxylipin 2E,4E-decadienal (DD) (Vardi *et al*., 2006; Vardi, 2008; Graff Van Creveld *et al*., 2021). Cell death measurements 24 hours post infochemical treatment did not reveal significant differences in the response of the triple MC KO and the WT strain (Fig S7). Together, these results strongly suggest that MCs have specific protective roles during thermal stress that is not a general stress response.

**Figure 4.**
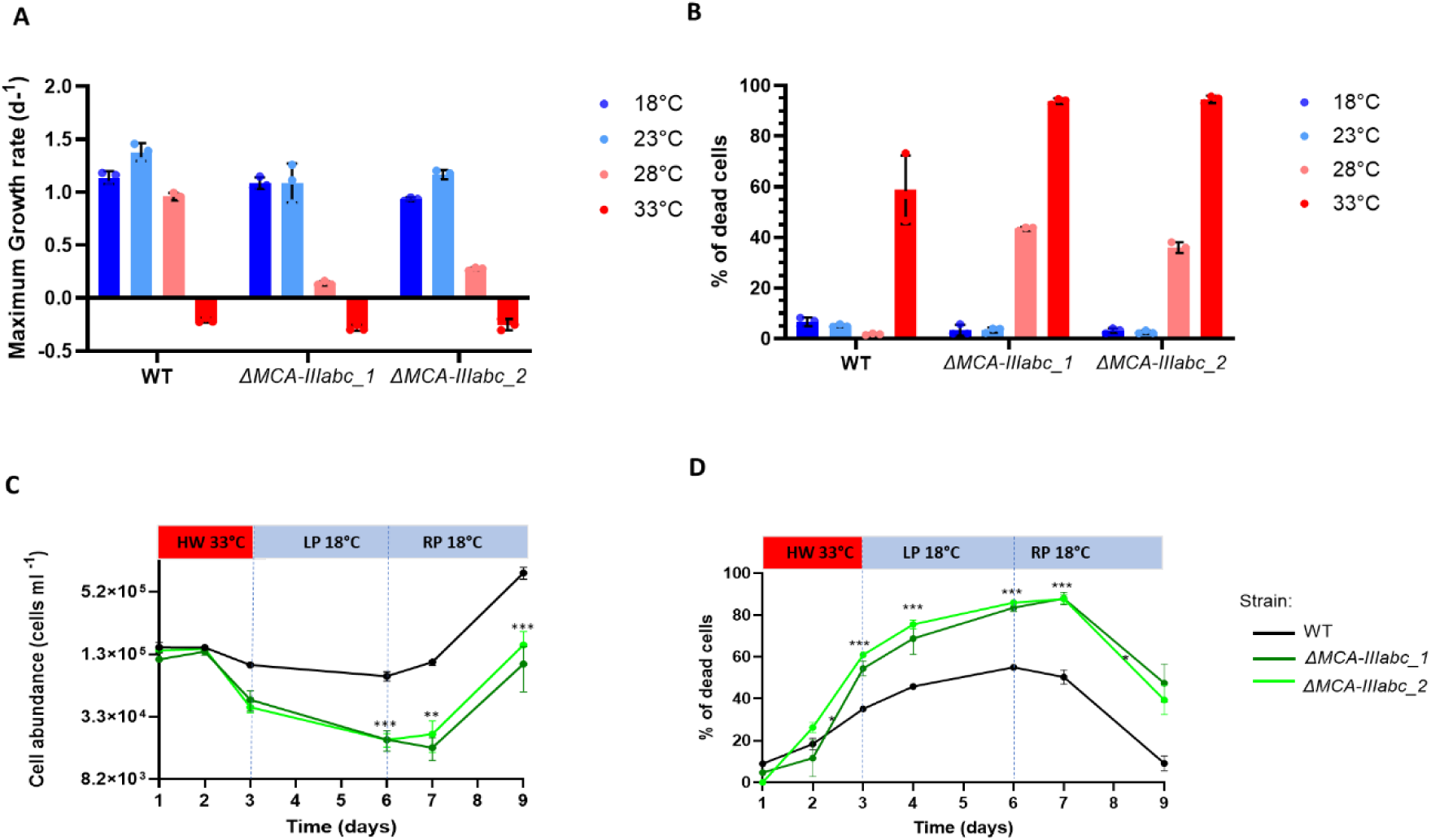
Triple MC KO lines are hyper-sensitive to elevated temperature and heatwave treatment. **(A)** Maximum growth rate of WT and two triple MC mutant strains in different growth temperatures. **(B)** Quantification of cell death in WT and two MC strains, measured by Sytox Green staining using flow cytometry after 5 days of growth in 4 different temperatures. **(C-D)** WT (black) and triple MC mutants (dark and light green) were exposed to 33°C for 72h and moved back to 18°C. Cultures were sampled for cell counts **(C)** and cell death **(D).** Values represent mean ± standard deviation, n=3. Statistical significance was calculated by repeated measures ANOVA as compared to WT. Asterisks represent P-values: * ≤ 0.05, ** ≤0.003, *** ≤ 0.001.

PtMCA-IIIc was originally identified in response to oxidative stress using redox proteomics approach (Rosenwasser *et al*., 2014), therefore, we sought to examine the interplay between heat stress and ROS production. We quantified intracellular ROS accumulation during heat stress using an H_2_O_2_-sensetive fluorescent stain (BES-H_2_O_2_) (Sheyn *et al*., 2016). During the first 6.5 hours of heat-treatment, cells at 33°C generated about 1.6-fold higher H_2_O_2_ as compared those at 18°C (Fig. 5A). PtMCA-III triple KO lines displayed a similar accumulation of H_2_O_2_ as compared to WT but produced a higher basal level of H_2_O_2_ at 18°C, therefore eventually reaching higher fluorescence following heat treatment (Fig 4A and Fig. S7). The higher basal level of intracellular H_2_O_2_ in the MCs KO might contribute to their hyper-sensitive response to elevated temperature due to higher oxidative stress. Indeed, triple KO lines exhibited increased sensitivity to sub-lethal concentrations of H_2_O_2_ (80 µM), which led to 36% ±5 dead cells as compared to only 12.5% ±3 in WT cultures (Fig. 5B). To support this result, we sorted single cells of WT and triple KO lines into agar plates 3 h after H_2_O_2_ treatment and tested for colony formation (survival) after 14 days. In agreement with the percentage of dead cells (Fig. 5B), the triple KO lines had less colony forming cells compared to WT (∼20-25% in KO lines compare to ∼55% in WT) (Fig S9).

**Figure 5.**
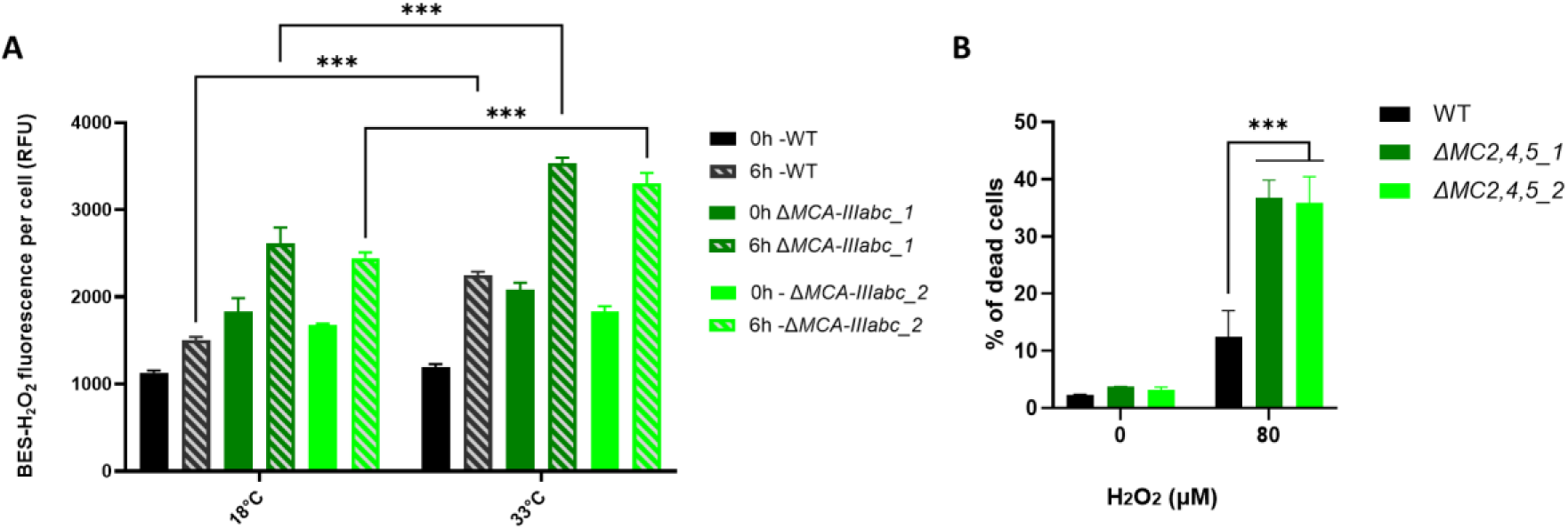
The response to heat stress is mediated by ROS accumulation that is enhanced in MC mutants. (**A**) Intracellular H_2_O_2_ levels as measured by mean fluorescence of BES-H_2_O_2_ detected by flow cytometry 0 and 6.5 hours after transition to 33°C or 18°C (control) in WT and MC triple KO mutants. (**B**) Cell death in MC triple mutants and WT populations 24 h after treatment with 80 µM H_2_O_2_ measured by Sytox staining. (**A-B**) Values represent the mean ± standard deviation, n=3. Statistical significance was calculated by one-way ANOVA compared to WT. * P-value ≤ 0.05, *** P-value < 0.0001.

## Discussion

Marine heat waves are likely to have major effects on phytoplankton blooms and especially on diatoms, a significant group in all global oceans that dominate regions prone to be affected by MHW (Feijão *et al*., 2018, 2020; Samuels *et al*., 2021; Rosalia Santoleri, 2024). Here, we characterized the thermal response of *P. tricornutum* and defined the distinct phases of acclimation by establishing a lab-based HW experimental system (Fig. 2, Fig. S1). By utilizing a functional genomics approach and generating MC mutants, we discovered a novel role for MC in mediating thermotolerance response.

*P. tricornutum* is highly sensitive to transient HW at a specific temperature, which led to suppression of its growth but induced only partial cell death in the entire population (50-60%), allowing survival of the rest of the population (Fig 2A-B). Interestingly, the peak of maximal cell death was observed 1-3 days after the HW had ended, when shifted back to the colder growth temperatures (Fig. 2B). Concomitantly, the heat-resistant subpopulation in *P. tricornutum* regains growth during the recovery phase (Fig 2A). These cellular dynamics in response to transient heat stress may imply phenotypic plasticity within the population and the establishment of co-existing resilient and susceptible subpopulations that can be part of population-level acclimation strategies to cope with elevated temperatures. Evidence of this cell-to-cell phenotypic variability has been shown previously in the response of *P. tricornutum* to high light and oxidative stress whereby survival of the population was improved by creating a sub-population resistant to stress that continues to grow (Mizrachi *et al*., 2019). An alternative explanation to the increase in cell death after returning to the optimal growth temperature can be the sensitivity of the heat-induced cellular machinery to cooling. Abrupt temperature effects were shown to cause stress, growth arrest and pigmentation production in *P. tricornutum* by causing lower carbon uptake (Rehder *et al*., 2023). Similarly, arctic phytoplankton communities also showed that regarding temperature oscillations, physiological responses to temperature shifts are less predictable than stable warming. While growth rates were not affected when temperatures rose, they were decreased during the cooling phase (Wolf *et al*., 2024). In addition, even following short heat-stress treatment, the returning to steady state cellular conditions is not a rapid transition. In *C. reinhardtii* cells exposed to 24 hours of temperature elevation from 25°C to 42°C, the protein concentration and cell diameter increased continued to rise up to 8 hours after the heat-stress was over (Hemme *et al*., 2014). In the current HW setup, it is still unclear whether returning to initial temperatures causes additional stress to the cultures or that phenotypic heterogeneity between the cells led to a delayed response in the cellular cascade leading to PCD.

### The vital role of metacaspases in thermotolerance

The molecular and biochemical drivers of acclimation to heat waves in marine diatoms are still unclear. In this work, we revealed a novel role for MCs in the thermotolerance of diatoms. MCs were classically associated with phytoplankton response to diverse environmental stress conditions, including iron limitation (Bidle & Bender, 2008), and oxidative stress (Graff Van Creveld *et al*., 2021) which ultimately led to induction of cell death.

Here we reveal a novel vital role of MCs in marine phytoplankton response to thermal stress. The three PtMCA-III genes show different levels of expression in replete medium, nitrogen and phosphate depletion and iron limitation (20, 40 and 400 pM Fe) (Smith *et al*., 2016; Matthijs *et al*., 2017; Graff Van Creveld *et al*., 2021). We demonstrated that KO lines of *P. tricornutum* that lack all three expressed PtMCA’s are highly sensitive to heat stress and are impaired in various physiological measures such as growth, photosynthetic activity and cell death (Fig 4). The hypersensitive response to heat was detected throughout the latent and recovery phases (Fig 4 C-D), implying that MC may also be necessary for the returning to initial cell state. KO of all three MC genes also caused a shift in the critical temperature that causes induction of cell death, demonstrating its pivotal role in determining thermotolerance regulation (Fig 4 A-B). We also show that the sensitivity of the triple KO is a specific response to elevated temperatures (Fig S7). The increased sensitivity to heat stress is in agreement with results observed in MC mutants in plants, yeast and in the green alga *C. reinhardtii* (Lee *et al*., 2010; Zou et al., 2023; Ruiz-Solaní *et al*., 2023). Thus, MC may have a conserved role in thermotolerance across eukaryotic unicellular organisms and plants from distant evolutionary origin. Since single and double KO lines do not show cell death or growth arrest in response to heat but have slower growth rates in steady state, we suggest that each MC might have a vital but different role. Similarly to *A. thaliana,* where some of the MCs have a role related to cell death, while others in clearing protein aggregates and delayed senescence (Coll *et al*., 2010; Watanabe & Lam, 2011; Ruiz-Solaní *et al*., 2023). Gene expansion and diversification of MCs, as described in *A. thaliana*, may have resulted in multiple cellular functions. A similar multitude of functions might act in diatom MCs whereby some are mediators of cell death and others, as shown here, have a vital role in acclimation and resilience to stressors such as elevated temperature. This is opposed to yeast and *C. reinhardtii* that have only a single MC gene, which was shown to have multiple roles involving cell homeostasis, senesces, cell death, and thermotolerance, depending on the physiological and cellular context (Lee *et al*., 2010; Hill *et al*., 2014; Zou et al., 2023). Recently, the involvement of MC in thermotolerance was also shown in *C. reinhardtii* where re-localization of the MC was observed following heat stress, concomitant with induction of its non-proteolytic role in modulating membrane permeability that prevents membrane fluidity in response to heat (Zou et al., 2023). Interestingly, the yeast MC Yca1, was shown to possess a dual function; a proteolytic role during cell death and chaperone-like activity during heat stress. The shift between these distinct cellular functions depends on calmodulin binding that inhibits the proteolytic activity and enables the chaperone-like activity (Eisele-Bürger *et al*., 2023). This versatility enables tight regulation of proteins associated with cell fate regulation, enabling tuning to the physiological status of the cells. Furthermore, Yca1 mutants displayed more protein aggregation and autophagic bodies after heat stress compared to WT (Lee *et al*., 2010). Thus, Yca1 was suggested to be involved in the clearance of protein aggregates during heat stress.

### Diatom’s cellular response to heat stress is mediated by H_2_O_2_ generation

In marine algae, various environmental stress conditions can cause ROS accumulation in the cell which can lead to the oxidation of lipids, nucleic acids and proteins (Vardi *et al*., 1999; Rosenwasser *et al*., 2014; Huang *et al*., 2025). At lower concentrations, ROS are crucial for signaling and homeostasis of the cell, while higher levels can lead to cell death (Mittler, 2017). The link between intracellular ROS elevation and heat stress exposure is highly studied in plants. ROS waves were associated with heat stress in *A. thaliana* and the production of ROS by mesophyll cells was required for acclimation to heat (Zandalinas & Mittler, 2021). In heat-treated plants, Rubisco activity is reduced leading to the production of H_2_O_2_ (Sharkey, 2005). In addition, ROS elevation during heat stress was associated with compromised oxygen-evolving complex (OEC) and inhibition of the PSII repair systems (Allakhverdiev *et al*., 2008; Song *et al*., 2014).

Heat treatment and ROS generation were shown in coral symbionts, dinoflagellates. Dinoflagellates acclimated to 25°C and transferred to 32°C significantly increased production of ROS molecules, H_2_O_2_ and superoxide anion (O_2_ ^-^) (Rosic *et al*., 2024). In the coral symbiont *Symbiodinium*, incubation at 33°C resulted in decline of maximum photosynthetic efficiency and elevation of superoxide dismutase (SOD) activity (Krueger *et al*., 2014). In our setup for diatom response to heat waves, photosynthesis shuts down completely as values of Fv/Fm reach almost zero, indicating the severe damage to PSII is possibly enhanced by ROS elevation (Fig 2C). Our results link heat stress and H_2_O_2_ production and reveal elevation of intracellular H_2_O_2_ levels during the first hours of the heat stress (Fig.5A). In addition, at least one of the MCs, PtMCA-IIIc, is redox-sensitive, allowing fine tuning of its activity and regulation of cell fate as a function of the extent and duration of stress conditions mediated by ROS signaling (Graff Van Creveld *et al*., 2021). The PtMCA-III mutants have higher intracellular H_2_O_2_ levels in steady state conditions. When treated with exogenous H_2_O_2_, they exhibited hypersensitivity and higher cell death as compared to WT cells (Fig. 5B). Similar effect of H_2_O_2_ sensitivity was also demonstrated in yeast Yca1 mutant strain that accumulated more carbonylated proteins than WT post H_2_O_2_ treatment (Khan *et al*., 2005). Here, the intracellular H_2_O_2_ concentration in both WT and triple mutant lines was elevated in response to heat, with almost the same fold change, suggesting that during heat stress ROS activates pathways that are upstream to MC (Fig 5A). Therefore, we suggest that perception of heat stress is mediated by induction in intracellular ROS which will activate a metacaspase-dependent signaling cascade, ultimately regulating cell fate.

In conclusion, this work provides novel insights into the unknown machinery of short-term exposure to heat stress and a novel vital role for metacaspases in mediating the response of diatom cells to thermotolerance. Future research is needed to deepen our understanding of the molecular components driving algal acclimation to heat-stress (Huang *et al*., 2025). Considering climate change and MHW spatiotemporal expansion, it is crucial to study how it will shape and impact diatom ecophysiology and biogeochemistry in contemporary and future oceans.

## Methods

### Culture maintenance

*P. tricornutum* culture, accession Pt1 8.6 (CCMP2561) was purchased from the National Center of Marine Algae and Microbiota (NCMA, formerly known as CCMP). Cultures were grown in filtered sea water (FSW) supplemented with f/2 media (Guillard & Ryther, 1962) at 18°C with 16:8 hours light:dark cycle and light intensity of 170 μmol photons m^-2^ sec^-1^ supplied by cool-white LED lights (Edison, New Taipei, Taiwan). Unless otherwise specified, all experiments were performed with exponentially growing cultures (∼5·10**^5^**-1·10^6^ cell ml^-1^ on the day of the experiment). For measuring growth rates, cultures were diluted to 1·10**^5^** cells ml^-1^ on day 0 of the experiment. Heatwave incubation conditions were with the same light patterns and intensities.

### Enumeration of algal cell abundance, cell death and intracellular ROS levels

Algal cells were quantified using the CytoFLEX flow cytometer (Beckman coulter, Indianapolis, USA). The cell population was identified by plotting the chlorophyll fluorescence (ex: 488 nm, em: 663 - 737 nm) versus forward scattered light (a proxy for cell size). At least 10,000 cells were sampled. Cell death analysis was done by Sytox Green (Invitrogen) staining, a nucleic acid stain that penetrates compromised membranes, which is a characteristic of dead or dying cells. The Sytox green stock solution was diluted to 5 mM in dimethyl sulfoxide (DMSO) and added to the samples at a final concentration of 1 μM. Then, samples were incubated in the dark for 30 minutes at room temperature and analyzed by flow cytometry using the green channel (ex: 488 nm, em: 525 nm). Intracellular H_2_O_2_ levels were measured with BES-H_2_O_2_ (Waco), a highly selective probe for cell-derived H_2_O_2_. Samples were stained with BES-H_2_O_2_ at a final concentration of 50 μM and then analyzed by flow cytometry (ex: 488 nm, em: 500-550 nm). Unstained samples were used as a control for each treatment, to eliminate the background signal. Growth rate was measured using flow cytometry in the exponential growth phase from day 2 to 5 according to the following equation (equation A): 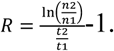

### Quantification of photosynthetic efficiency

Photosynthetic efficiency was measured using Water-PAM (WALTZ). Photosynthetic efficiency was determined as Fv/Fm, calculated as previously described (Maxwell & Johnson, 2000). Fm represents the maximum fluorescence emission level in the dark measured with a saturating pulse of light (emission peak at 450 nm, 2,700 μmol photons m^-2^ sec^-1^, 800 ms); Fv = Fm – F0.

### Generation of metacaspases *P. tricornutum* mutants using CRISPR/Cas9

For MC KO, we utilized previously constructed *ΔMCA-IIIc* cell line where the MC5 (ID:Phatr3_J54873.t1) was knocked out and cells express the Cas9 gene (Graff Van Creveld *et al*., 2021) and a protein that confers resistance to Zeocin. To this cell line we introduced 4 sgRNAs on one plasmid to create a double KO of PtMCA-IIIb (ID: Phatr3_J54872.t1) and PtMCA-IIIc (*ΔMCA-IIIbc*) and triple KO for PtMCA-IIIa (ID: Phatr3_J48151.t1), PtMCA-IIIb and PtMCA-IIIc (*ΔMCA-IIIabc_1*) (Fig S2). Transformation was achieved by biolistic particle delivery system (Bio-Rad), as previously described (Apt *et al*., 2002). Briefly, cells were plated on agar plates 2 days prior to bombardment at a concentration of 2×10^7^ cell per plate (50% FSW+F/2, 1.5% agar, without antibiotics). M17 W-particles (Bio-Rad) were coated with 1 μg/μl DNA in the presence of CaCl_2_ (2.5M) and spermidine (0.1M). Cells were bombarded using a rupture disc of 1300psi, using the hepta-adaptor according to the manufacturer’s instructions. Following the biolistic shooting, cells were transferred to recover on selection plates (50% FSW+F/2, 1.5% agar with 100 mg/ml Nourseothricin). Colonies that grew on selection plates were isolated and sub-cloned on plates with 100 μg/ml Zeocin (InvivoGen) and Nourseothricin (Jena Bioscience). To verify the CRISPR-Cas9 knockout for all cell lines, cell lysates of antibiotic-resistant colonies were prepared in lysis buffer (1% TritonX-100, 20 mM Tris–HCl pH 8, 2 mM EDTA) and subjected to repeated freezing and thawing. Then, cell lysate (5 μl) was used for PCR amplification of the genomic targets with primers specific to the gene of interest. To confirm that the mutagenesis has caused frameshifts, insertions or deletions, the target regions were amplified using specific primers designed to flank the targets. Random clones were sent to Sanger sequencing to validate deletion. Analysis was done using the sequence alignment programs Sequencher and Snapgene.

### H_2_O_2_, DD and BrCN treatments

*P. tricornutum* cells were treated with H_2_O_2_ in 24 well plates (2 ml culture per well) and monitored for cell death as described above. H_2_O_2_ (diluted in DDW) was added to the cultures to the final concentration of 80 μM. Stock solution of 95% DD (Acros Organics) was diluted in MeOH and added to the cultured at 25, 50 and 100 μM. Control cultures were treated with 0.25% of MeOH. BrCN (Sigma) treatment was induced with three doses (1.75, 2.5, 5 μM), and control cells were treated with 0.1% acetone. After treatment, cells were incubated for 24 hours under normal growth conditions and analyzed for cell death as described above.

### Single cell sorting after stress treatment

WT *P. tricornutum* and triple MC mutant cells were treated with H_2_O_2_ (80, 100 and 120 μM) for 3 hours, then sorted using BD FACS ARIA III (New Jersey, USA). Single cells were sorted directly on one well agar plate (FSW+f/2, 1.5% agar). Agar plates were screened for colony formation two weeks after sorting by the Typhoon™ Laser Scanner (Cytiva, Marlborough, Massachusetts, USA) using the Cy5 laser.

### Metacaspase proteolytic activity assay

Kinetic measurement of MC activity was performed *in vitro*. A total of ∼1×10^8^ cells were harvested at the exponential growth phase (∼1×10^6^ cells ml^-1^) by centrifugation and plunged into liquid nitrogen. Then, cell pellets were resuspended in 250 μl lysis buffer (150 mM NaCl, 25 mM HEPES, 10% glycerol, 0.2% triton, 1 mg ml^-1^ lysozyme, 1 μl benzonase, 0.5 mM DTT, pH 7.8) and sonicated (10 sec of sonication followed by 10 sec of rest, 10 cycles). For activity assays, protein concentration was determined by the BCA method and the sample concentration was normalized in base buffer (150 mM NaCl, 25 mM HEPES, 10% glycerol). Assays were performed in activity buffer (base buffer with 0.1% CHAPS, 10 mM DTT, 10 mM CaCl_2_). VRPR (Ac-Val-Arg-Pro-Arg) conjugated to the fluorophore 7-Amino-4-methylcoumarin (AMC, NBT-Bachem) was used as the substrate. As control, the MC inhibitor VRPR-fmk (NBT, final concentration 25 μM), was incubated in activity buffer with the cell lysate for 30 min before substrate addition. Trypsin was used as a positive control, as this protease cleaves after arginine and lysine. The fluorophore AMC was used for standard curve (12, 6, 3, 1.5 and 0.75 μM AMC). Fluorescence was measured using the Infinite 200 pro, Tecan (Männedorf, Switzerland) plate reader, with ex: 360 nm and em: 460 at 20°C. The fluorescence measurement started 1 min after substrates were added to the lysates, for 90 minutes with a 1-minute interval. The activity rate was calculated after extracting the maximum slope (RFU/min), multiplying by the AMC standard slope, and normalizing according to protein concentration (μmol AMC·min^-1^·mg protein^-1^).

## Acknowledgments

This research was supported by the Israeli Science Foundation (ISF) (grant # <u>749/24</u>) awarded to A.V. and D.S. and by the Tom and Mary Beck Center for Renewable Energy as part of the Institute for Environmental Sustainability (IES) at the Weizmann Institute of Science. MS was also supported by the Institute for Environmental Sustainability (IES) at the Weizmann Institute of Science.

## Author contributions

MS and AV planned and designed the research, analyzed the data and wrote the article. SG designed the plasmids for mutant transformation. AM, AZ, DS and MS carried out the mutant transformation. SBD and MS performed the mutant sequence analysis and homozygous validation.

## Supporting information

### Supplementary Figures

**Supplementary Figure S1.**
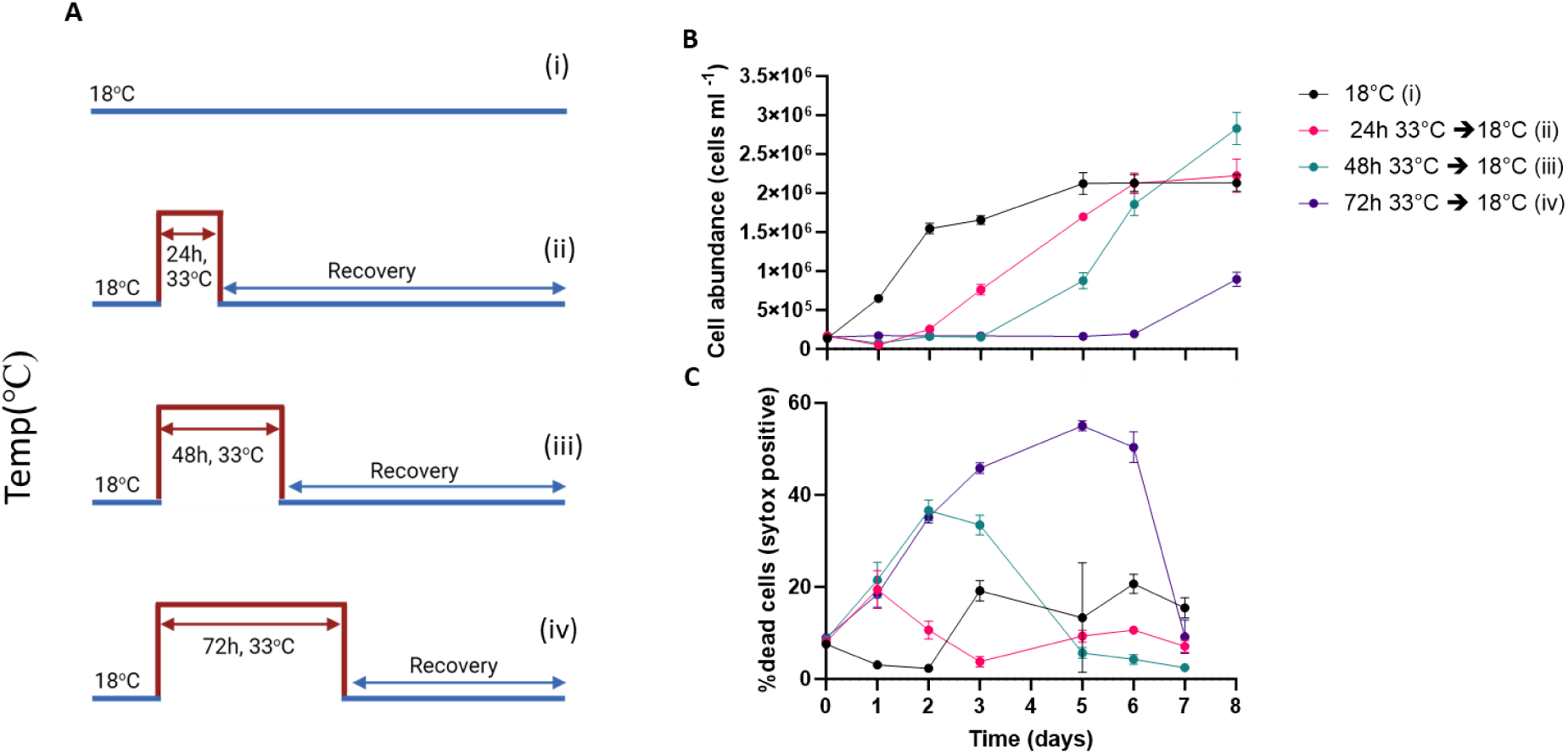
Experimental setup for heat wave treatment at different intervals. (**A-C**) Cultures grown at 18° C were exposed at day 0 to heat wave (HW) treatments of 33°C for 24h (ii, pink), 48h (iii, cyan) or 72h (iv, purple), and then transferred back to 18°C for recovery. Control cultures (i, black) remain for the duration of the experiment at 18°C. (A) Schematic representation of experimental design for HW duration calibrations. Cells abundance (B) and percent dead cells (C) over time in cultures treated as described in (A).

**Supplementary Figure S2.**
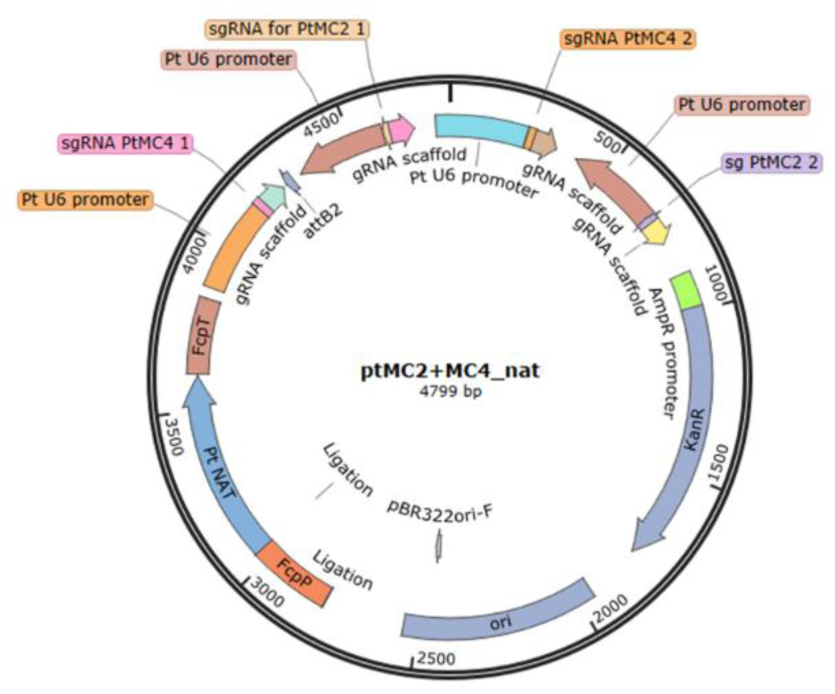
Plasmid design for PtMCA-IIIb and PtMCA-IIIc KO in *P. tricornutum*. The plasmid consists of a U6 promotor and a Nourseothricin (NAT) resistance gene. In addition, there are 2 sets of sgRNA. Background cell line for MC already has the Cas-9 therefore there is no Cas-9 in the plasmid.

**Supplementary Figure S3.**
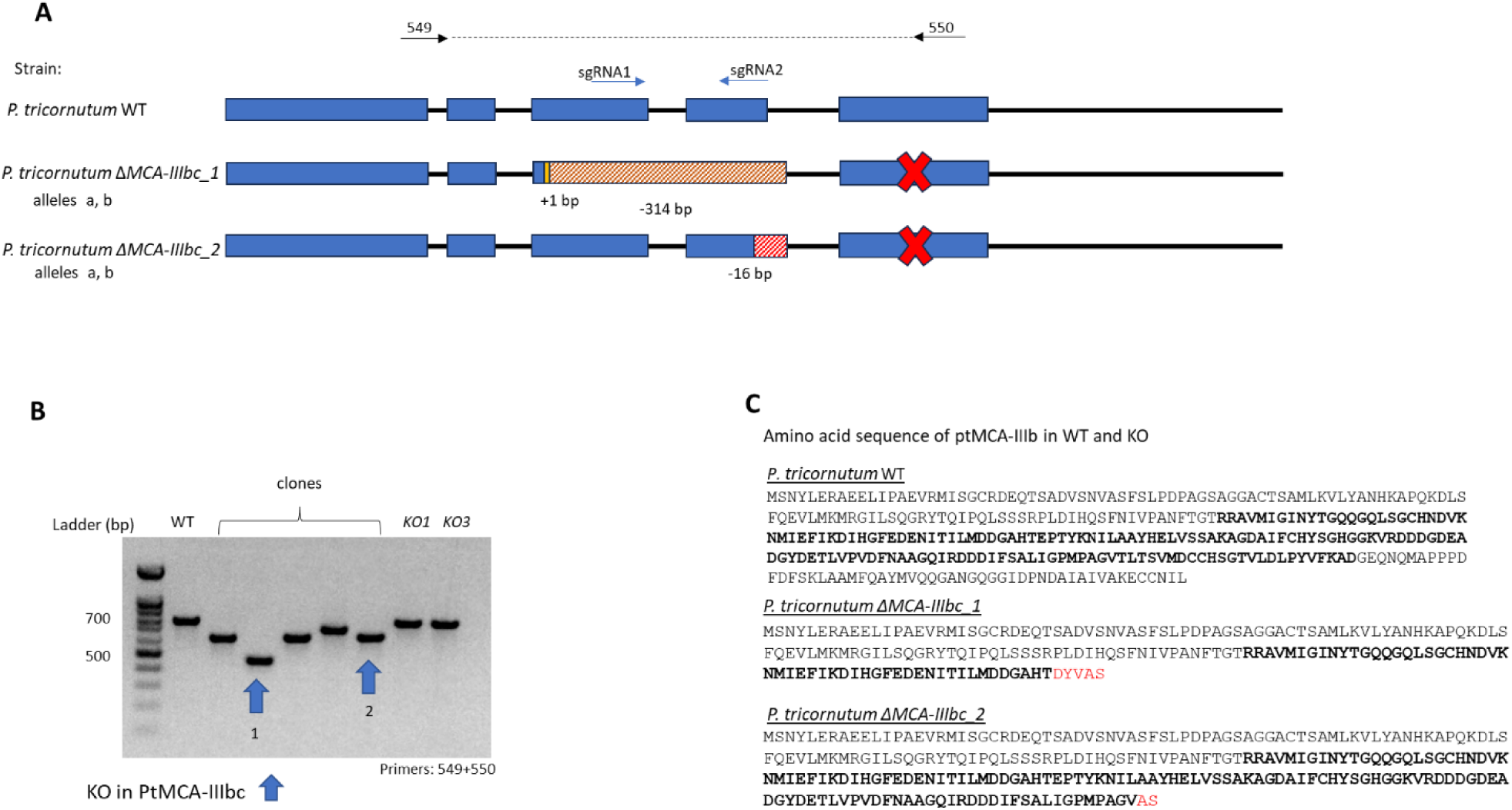
Creating MC-double mutants by knockout of PtMCA-IIIb in *P. tricornutum* ΔPtMCA-IIIc cells. **A.** Schematic representation of WT and the two clones with a deletion in the PtMCA-IIIb gene. Blue area represents the 5 exons, black lines are the introns. Red dashed rectangles represent deletions. In clone *ΔMCA-IIIbc_1* there is a 1 bp insertion of thymine before the deletion of 314 bp that is also partly in the junction therefore there is no 5^th^ exon (marked with a red X). Clone *ΔMCA-IIIbc_2* has a 16 bp deletion, 10 bp are on the 4^th^ exon and 6 bp are on the junction therefore there is no splicing and no 5^th^ exon. In both lines both alleles are identical. Blue arrows are in the location of the two guide RNA, black arrows are the primer location used for validation. **B.** Verification of deletion in the PtMCA-IIIb DNA sequence by PCR amplification with 549+550 primers (as depicted in A). Blue arrows designate the two chosen mutants with the 314 and 16 bp deletions. KO1 and KO3 are the single KO in PtMCA-IIIc. **C**. Amino acid sequence of the PtMCA-IIIb gene. Bold letters represent the p20 domain. *ΔMCA-IIIbc_1* has a frame shift causing a replacement of 155 amino acids with only 5 novel ones, followed by a stop codon. *ΔMCA-IIIbc_2* has a frame shift causing a replacement of 77 amino acids with only 2 different ones, followed by a stop codon (in red).

**Supplementary Figure S4.**
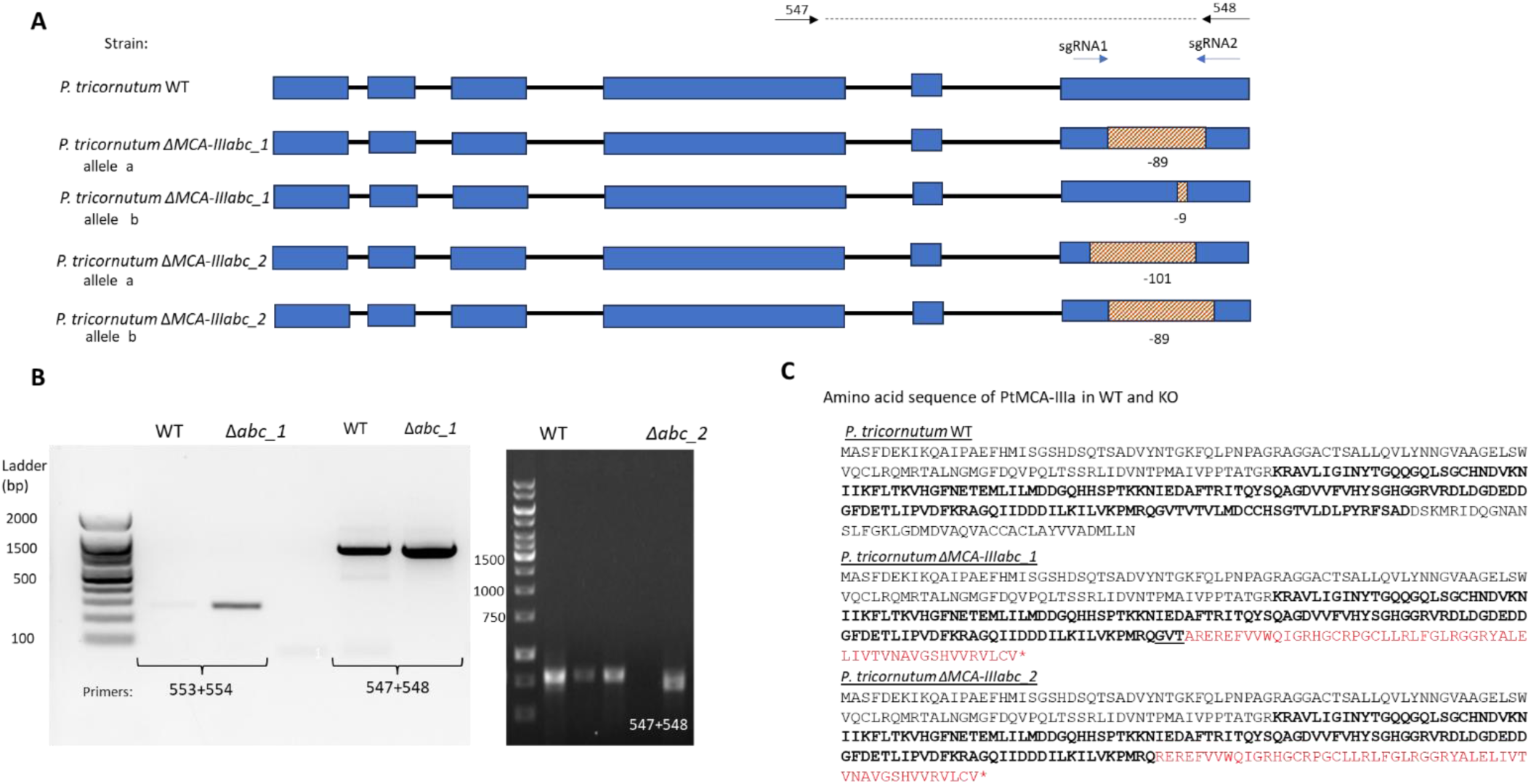
Generation of MC-triple mutants by knockout PtMCA-IIIa and PtMCA-IIIb in *P. tricornutum ΔMCA-IIIabc* cells. **A.** Schematic representation of the two clones with a deletion and WT DNA sequence of PtMCA-IIIa gene. Blue area represents the 6 exons, black lines are the introns. Red dashed rectangles represent deletions in each allele. Blue arrows are in the location of the two guide RNAs, black arrows are the primers’ location used for validation **B.** Verification of deletion in the PtMCA-IIIa DNA sequence by PCR amplification with 547+548 primers for the 9 bp deletion and 553+554 primers that are at the junction of the 89 bp deletion so result only in a band for the mutant. **C**. Amino acid sequence of the PtMCA-IIIa gene. In bold is the p20 domain. Red letters represent novel amino acids added because of the DNA frame shift. *ΔMCA-IIIabc_1* has a 65 amino acid deletion and 53 novel amino acids on one allele and a 3 amino acid deletion on the other allele (underlined). *ΔMCA-IIIabc_2* has a deletion of 68 amino acid deletion and an introduction of 52 novel amino acids and no stop codon (in red). Both alleles have the same frame shift.

**Table S1.**
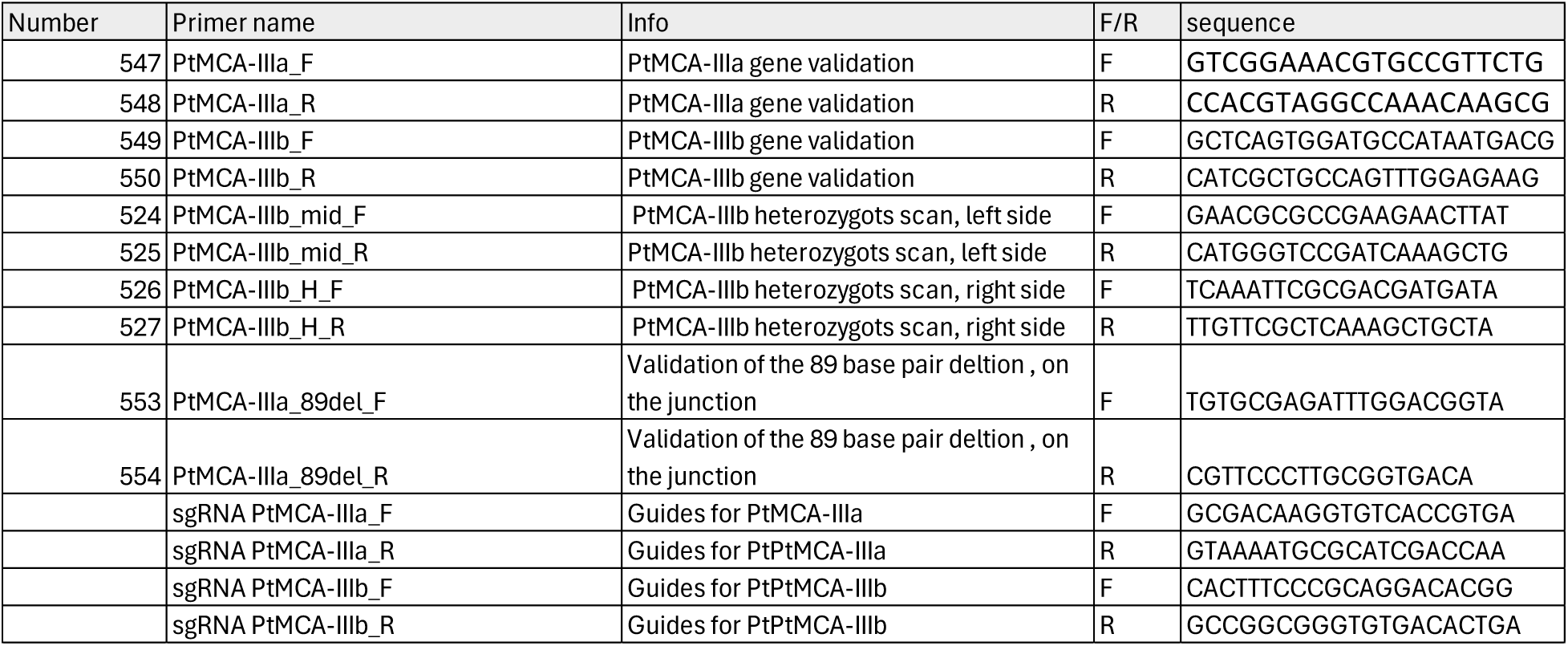
Guide RNA and primer sequences used in MC transformation.

**Supplementary Figure S5.**
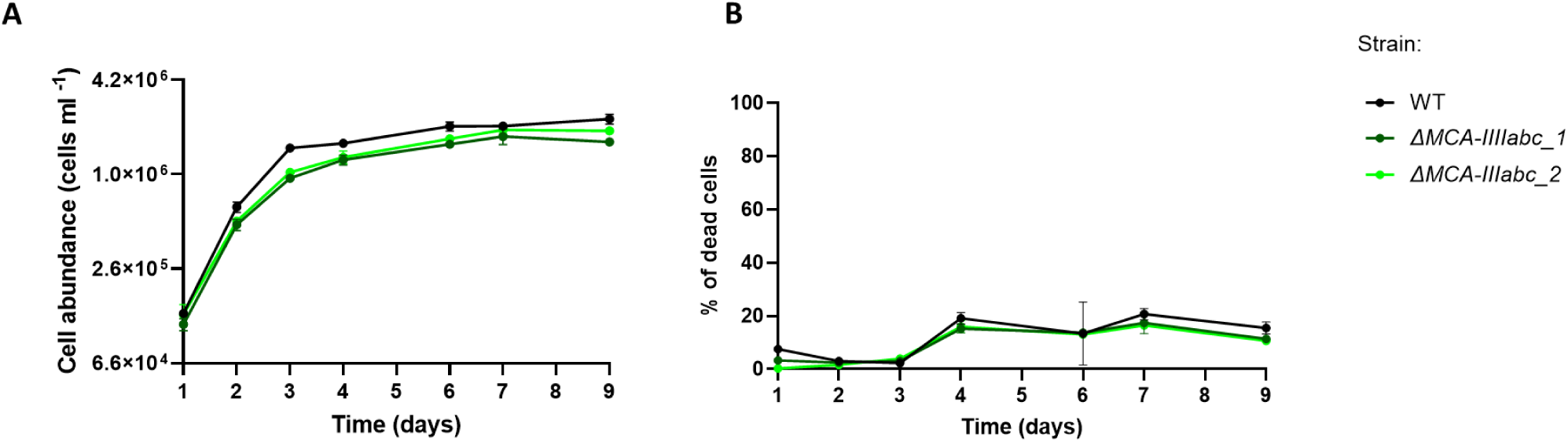
Growth curves of WT and triple MC KO under normal conditions. Cell counts (A) and cell death (B) of WT (black) and triple MC KO (greens) grown in 18℃. Values represent the mean ± standard deviation, n=3.

**Supplementary Figure S6.**
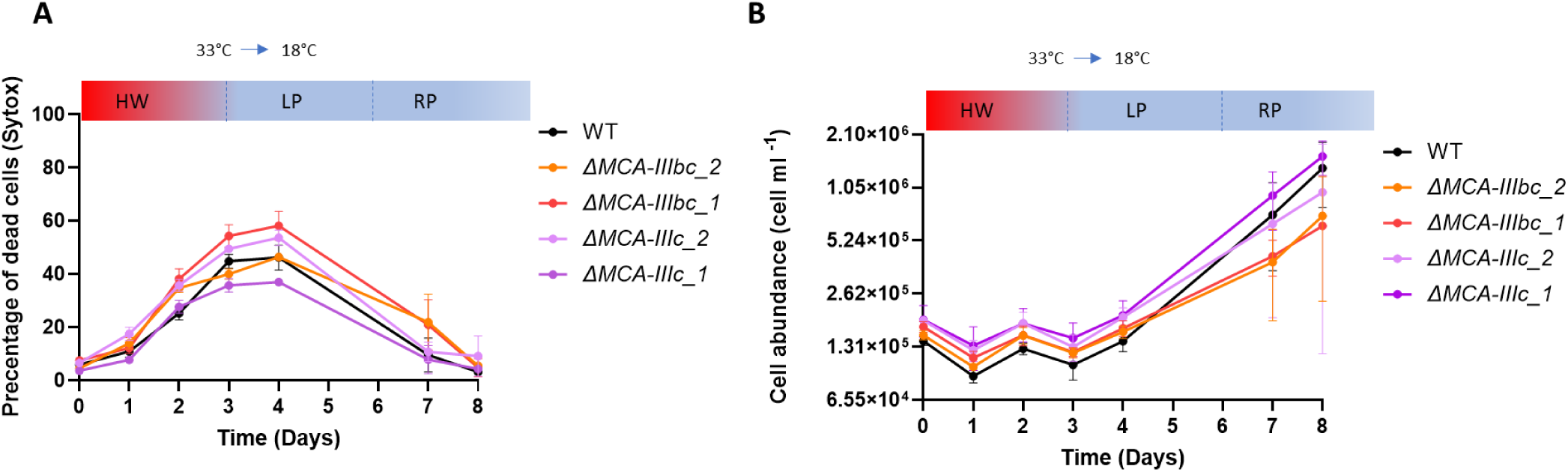
HW treatment of WT, single and double MC mutant lines. Quantification of cell death (A) and cell counts (B) of WT (black), single (in orange and red) and double (light and dark purple) mutants exposed to 33℃ for 72 hours and transferred back to 18℃ (black line). Values represent the mean ± standard deviation, n=3.

**Supplementary Figure S7.**
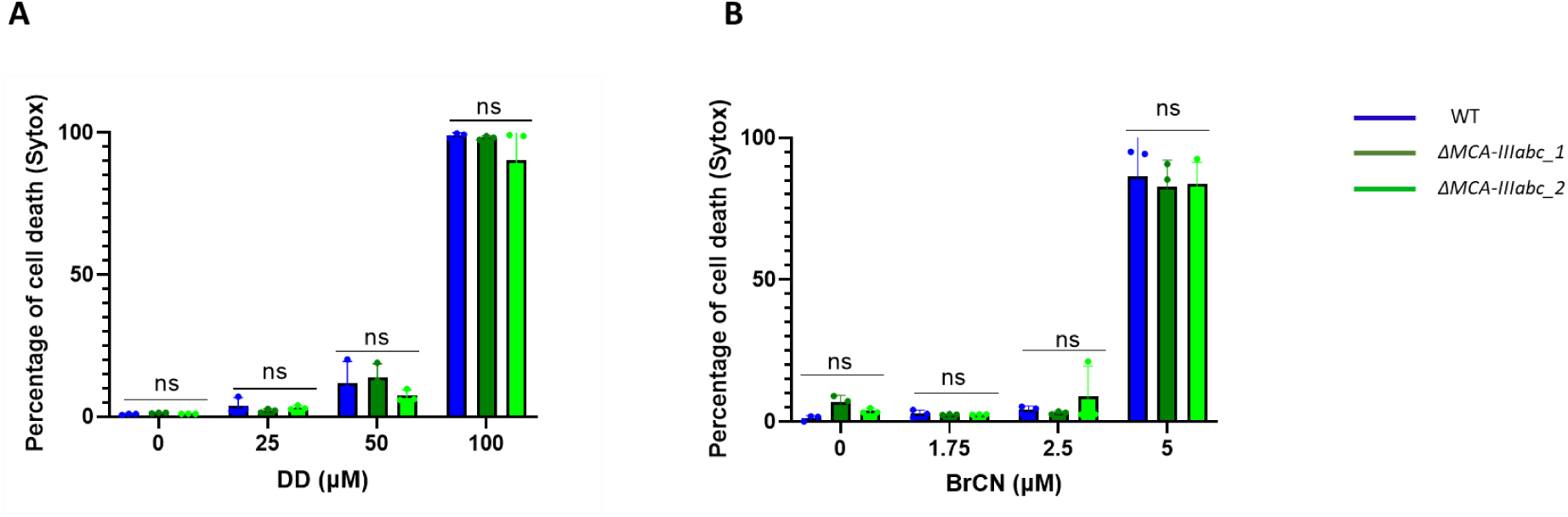
Triple MC KO are not hypersensitive to DD and BrCN treatments. **A**. Percent of Sytox positive cells in cultures treated with 25, 50 and 100 μM of DD. **B**. Percent of Sytox positive cells in cultures treated with 1.75, 2.5 and 5 μM of BrCN. Statistical significance was calculated by using one-way ANOVA with *p* value < 0.05. Values represent the mean ± standard deviation, n=3. ns = not significant.

**Supplementary Figure S8.**
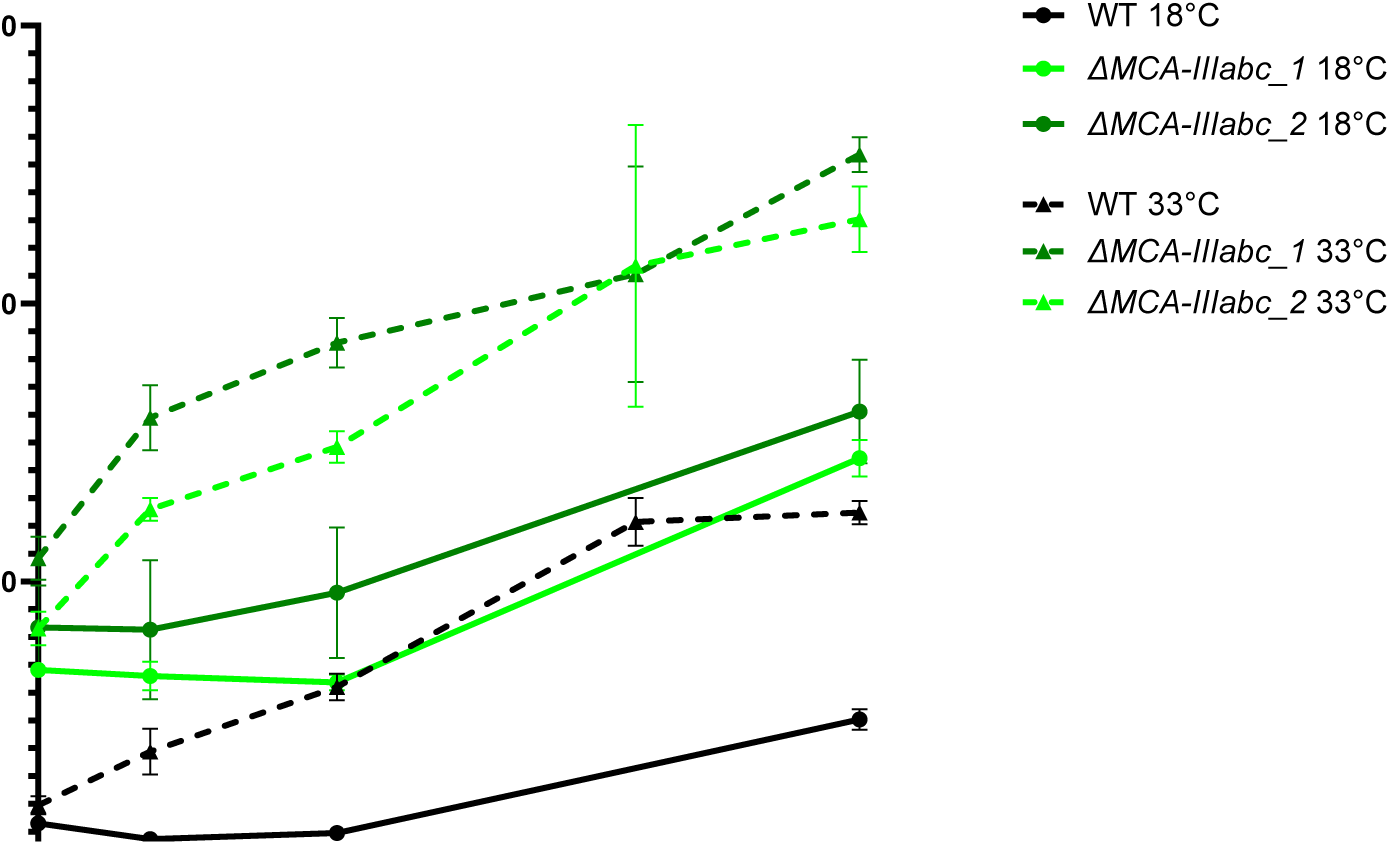
H_2_O_2_ production during exposure of diatom cells to heat stress. Mean fluorescence of WT and triple MC mutants stained with BES-H_2_O_2_ during incubation in 33°C for 6.5 hours. Solid lines are cultures grown at 18°C, dashed lines are cultures grown in 33°C. n=3.

**Supplementary Figure S9.**
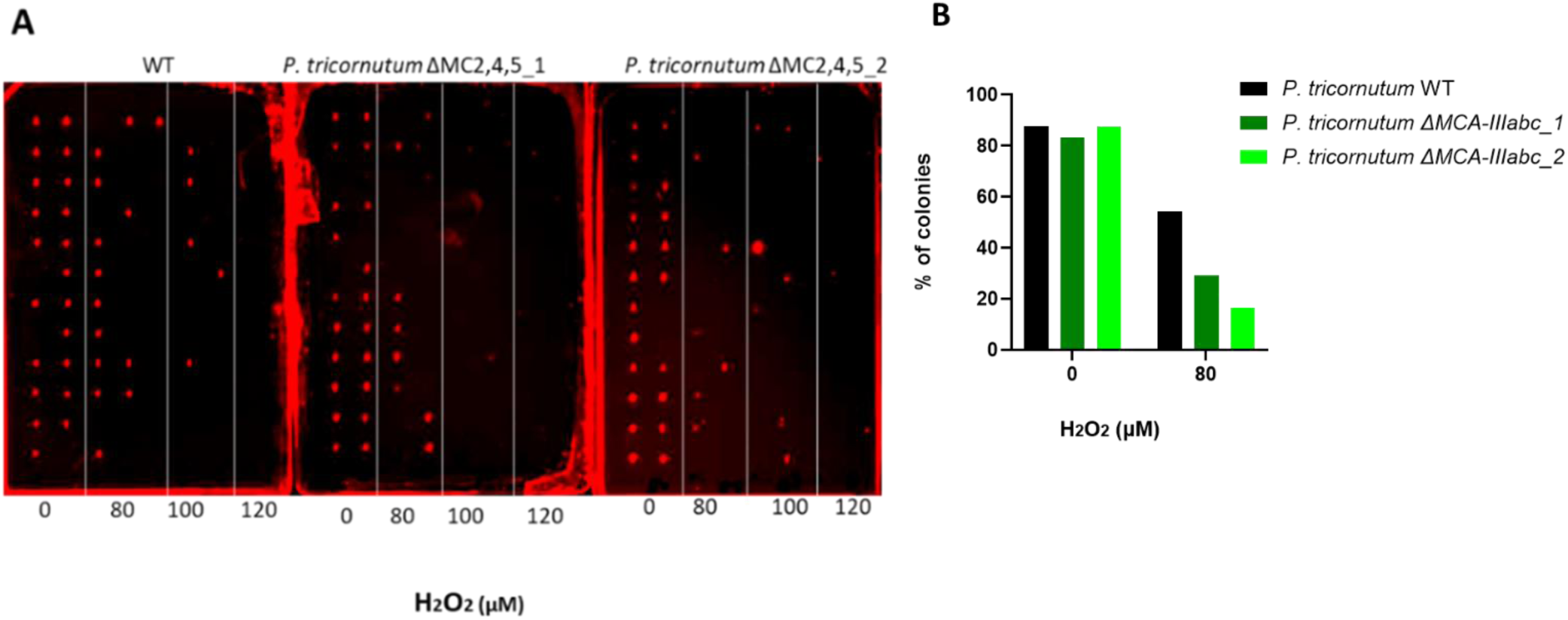
MC triple KO have higher sensitivity to oxidative stress. **A.** *P. tricornutum* mutant and WT cells were sorted onto agar plates (single cell per drop), 3 hours after treatment with 80, 100 or 120 µM H_2_O_2_. The plates were scanned for colony formation 14 days after sorting (see methods). **B.** Quantification of survival (colony growth) after 80 μM H_2_O_2_ treatment from a single experiment.

